# Dual site-specific chemoenzymatic antibody fragment conjugation using CRISPR-based hybridoma engineering

**DOI:** 10.1101/2020.12.08.415182

**Authors:** Camille M. Le Gall, Johan M.S. van der Schoot, Iván Ramos-Tomillero, Melek Parlak Khalily, Floris J. van Dalen, Zacharias Wijfjes, Liyan Smeding, Duco van Dalen, Anna Cammarata, Kimberly M. Bonger, Carl G. Figdor, Ferenc A. Scheeren, Martijn Verdoes

## Abstract

Functionalized antibodies and antibody fragments have found applications in the fields of biomedical imaging, theragnostics, and antibody-drug conjugates (ADC). Antibody functionalization is classically achieved by coupling payloads onto lysine or cysteine residues. However, such stochastic strategies typically lead to heterogenous products, bearing a varying number of payloads. This affects bioconjugate efficacy and stability, as well as its in vivo biodistribution, and therapeutic index, while potentially obstructing the binding sites and leading to off-target toxicity. In addition, therapeutic and theragnostic approaches benefit from the possibility to deliver more than one type of cargo to target cells, further challenging stochastic labelling strategies. Thus, bioconjugation methods to reproducibly obtain defined homogenous conjugates bearing multiple different cargo molecules, without compromising target affinity, are in demand. Here, we describe a straightforward CRISPR/Cas9-based strategy to rapidly engineer hybridoma cells to secrete Fab’ fragments bearing two distinct site-specific labelling motifs, which can be separately modified by two different sortase A mutants. We show that sequential genetic editing of the heavy chain (HC) and light chain (LC) loci enables the generation of a stable cell line that secretes a dual tagged Fab’ molecule (DTFab’), which can be easily isolated in high yields. To demonstrate feasibility, we functionalized the DTFab’ with two distinct cargos in a site-specific manner. This technology platform will be valuable in the development of multimodal imaging agents, theragnostics, and next-generation ADCs.

## II. Introduction

The use of antibody-drug conjugates (ADCs) has emerged as a potent strategy in the treatment of malignancies. As of late 2020, nine FDA-approved ADCs^1–9^ are used in the clinic, and several hundred are currently under clinical inestigation^10^. First- and second-generation ADCs are classically produced by conjugation of drug molecules to the side chains of solvent-exposed lysines or interchain cysteines.^11^ However, such approaches lead to highly heterogeneous end-products with variable molecular weights, drug coupling sites, and drug-to-antibody ratio (DAR), with the concomitant risk of influencing target binding affinity.^12^ Indeed, monoclonal antibodies (mAbs) typically contain more than 60 accessible lysines, whereas the drug-to-antibody ratio (DAR) should remain low enough (3-4) to prevent aggregation.^13,14^ Third-generation ADCs aim to address these challenges by using site-specific conjugation methods^11,12^. As opposed to random coupling, site-specific modification enables strict control over payload conjugation to generate a homogenous product.

Antigen-binding fragments (Fab’) are molecules derived from mAbs^15^. Their heavy chain (HC) is truncated to solely contain the variable domain VH and the constant domain CH1, enabling association with the light chain (LC), but lacks the CH2 and CH3 domains that dimerize to generate the Fc domain. While these Fab’ retain binding ability to their target, they do not exhibit Fc-mediated immune effector functions such as recruitment of effector cells, or fixation of complement^16^. Moreover, they have a shorter half-life in circulation^17,18^, and are more efficient at penetrating dense tissues in which conventional mAbs are excluded^17,19^. However, the chance of modifying the binding region of a Fab’ using classical stochastic labelling is higher than on full size mAbs, due to the smaller size and reduced number of reaction sites^20^. Thus, Fab’ fragments represent attractive proof-of-concept candidates for third-generation ADCs, as well as for imaging and theragnostic applications.

Functionalization of antibody fragments with distinct payloads is an attractive strategy in for several applications. While combination therapies are gaining more attention in chemotherapeutic treatments, classical ADCs target only one drug to cancer cells. Similarly, multimodal imaging enables the visualization of targets of interest on different scales, from whole body imaging with radioisotopes down to the histological level with fluorescent tracer molecules. These applications would benefit from the development of a flexible plug-and-play antibody fragment engineering platform for dual site-specific labelling. Most-site specific conjugation strategies make use of a short peptide tag (e.g. a sortase A recognition motif^21^), or engineered residues^11,22^ to introduce cargos. Thus, they only permit functionalization with multiple distinct payloads through the synthesis of orthogonal multivalent linker systems or multifunctional conjugates, with concomitant synthetic and potential solubility issues. Here, we report a widely applicable strategy to introduce two orthogonal site-selective labelling tags on a Fab’ fragment by capitalizing on our recently reported Clustered Regularly Interspaced Short Palindromic Repeats/Homology Directed Repair (CRISPR/HDR) hybridoma genomic engineering approach^23^. In this work, we expand the genomic engineering toolbox to enable modification of the HC and LC loci of the mouse IgG1 (mIgG1) hybridoma, available for a plethora of targets. With this, dual-tagged Fab’ (DTFab’) are generated equipped with two distinct sortase A recognition motifs (sortags) on the HC and LC, each orthogonally recognized by a specific variant of the “evolved sortase A” (eSrtA) enzyme (eSrt2A-9 or eSrt4S-9)^24^. These enzymes enable the ligation of virtually any payload bearing a synthetically easily accessible *N*-terminal polyglycine motif onto the target protein. To demonstrate feasibility, the DTFab’ were sequentially functionalized with two distinct cargos in a site-specific manner and thoroughly characterized. We expect that this technology platform will be a valuable asset in the development of Fab’ conjugates for imaging, theragnostic, and next-generation ADC applications.

## III. Materials and Methods

### Cell lines and general culture conditions

Human CD20-expressing BJAB cells and the mIgG1 anti-CD20-producing hybridoma clone (further referred to as anti-CD20 WT) were kept in RPMI1640 medium (11875093, ThermoFisher Scientific), 10% heat-inactivated foetal bovine serum (FBS), 2 mM Ultraglutamine-1 (BE17-605E/U1, Lonza), 1×Antibiotic-Antimycotic (15240062, ThermoFisher Scientific). Media was additionally supplemented with 50 μM 2-mercaptoethanol (2-ME) (M6250, Sigma Aldrich). Cells were kept at 37°C in a 5%CO_2_ humidified incubator and passaged three times per week. Cells were checked every five weeks for mycoplasma status.

### CRISPR/Cas9 plasmids generation

The genomic sequence of the Ighg1 constant heavy chain and Igkc constant light chain were identified using the Ensembl genome browser (release 98^25^, accession numbers – Ighg1: ENSMUSG00000076614, Igkc: ENSMUSG00000076609). gRNA-H.m1 (5’-CTTGGTGCTGCTGGCCGGGT -3’) and gRNA-L.mκ (5’-GGAATGAGTGTTAGAGACAA -3’) were designed using http://crispor.tefor.net/^26^ and ordered as single-stranded oligos from Integrated DNA Technologies (IDT) with the appropriate BbsI overhangs for cloning into the plasmid px330-U6-Chimeric_BB-CBh-hSpCas9, which was obtained as a gift from Feng Zhang (Addgene plasmid #42230). Oligos were phosphorylated using a thermocycler with the T4 PNK enzyme (M0201, New England Biolabs) by incubation at 37°C for 30 min, and annealed by incubation at 95°C for 5 min and cooling down at a rate of 0.1°C/sec to 25°C. Annealed oligos of gRNA-H.m1 and gRNA-L.mκ were cloned into the px330 vector at the BbsI site. Synthetic gene fragments containing homologous arms and desired insert for Fab′ fragment generation and tag insertion were synthesised by IDT and cloned into the PCR2.1 TOPO TA vector (K-450001, ThermoFisher Scientific). All gRNA/Cas9 and HDR plasmids were isolated from DH5α competent *E. coli* and purified with the NucleoBond Xtra Midi Kit (740410.100, Macherey-Nagel) according to the manufacturer’s protocol.

### Transfection

Nucleofection of the HDR template and CRISPR-Cas9 vectors was performed as described previously^23^. Briefly, hybridoma cells were assessed for viability, centrifuged (90 g, 5 min), resuspended in phosphate buffered saline (PBS)/1% FBS, and centrifuged again (90 g, 5 min). 1*10^6^ cells were resuspended in 100µL of SF medium with 1 μg of HDR template and 1 μg of Cas9 vector, or 2 μg of GFP vector (control) and transferred to cuvettes for nucleofection with the 4D-Nucleofection System (V4XC-2024, Lonza, CQ-104, Program SF). Transfected cells were quickly transferred to a six-well plate in 6 mL of prewarmed complete medium. The following day, the cells were transferred to a 10 cm petri dish in 10mL of complete medium supplemented with blasticidin (10 µg.mL^-1^, ant–bl-05, Invivogen) or puromycin (5 µg.mL^-1^, ant-pr-5, Invivogen), which were [preliminarily](https://m.linguee.fr/anglais-francais/traduction/preliminarily.html) titrated on the wild type hybridoma cells. Antibiotic pressure was sustained until GFP-transfected hybridomas were not viable, and HDR-transfected cells were confluent (typically between day 10 and 14 for blasticidin, and day 4 and 7 for puromycin). Subsequently, antibiotic-resistant cells were clonally expanded by seeding the hybridomas at the concentration of 0.3 cells per well in 100 μL of complete medium in U-bottom 96-well plates (limiting dilution). After 10 days, supernatant from wells with a high cell density was collected for further characterization.

### Genomic DNA isolation

One week after nucleofection with HDR and targeting constructs, a minimum of 1*10^4^ cells of the surviving cell population (bulk) were collected for genomic confirmation of HDR insertion. DNA was extracted using the Isolate II Genomic DNA kit (BIO-52067, Bioline), and was resuspended in ultrapurified water. The target region was PCR amplified using forward primers that anneal in the region directly upstream of the 5’HA for each locus, either fwd.m1 (5’-GTGCCGACTTCAATGTGCTT-3’) or fwd.mκ (5’-GTGCTTGTGTTCAGACTCCC-3’), and a reverse primer that anneals with the IRES in the HDR template rv.ires (5’-GGCTTCGGCCAGTAACGTTA-3’). PCR products were visualized on a 1% (w/v) agarose gel containing Nancy-520 dye (01494, Sigma-Aldrich).

### Flow cytometry

Ten days after limiting dilution, 50µL of supernatant from wells with high cell densities was transferred to wells in a V-bottom 96-well plate containing 5*10^4^ hCD20-expressing BJAB cells. After 20 min of incubation at 4°C, plates were centrifuged (300g, 2 min), and supernatant was discarded by flicking. Plates were washed twice with PBS/5% FBS and incubated with anti-is-tag (PE, 362603, BioLegend), anti-mIgG1 (PE, 406608, BioLegend) or anti Myc-tag (AF488, 2279, Cell Signalling Technology). To assess the binding capacity of constructs following CRISPR/Cas9 editing, and several steps of transpeptidation and purification, a competitive binding experiment was performed. Briefly, anti-CD20 parental antibody isolated from the untouched cell line supernatant was dialysed to borate buffer (pH=8.5) by ultracentrifugation, and 100 µg was incubated with 6 equivalents of NHS-AF647 (APC-005-1, Jena Bioscience, MW=1274 Da, 10 mM stock in DMSO) for 2h at RT in less than 40 µL. Excess label was quenched by diluting the reaction to a final volume of 50 µL using [50mM Tris pH=8.0], and incubating 15 min at RT. Labelled antibody was used without further purification. 5*10^4^ BJAB cells per well were seeded in a V-bottom 96-well plate and incubated in 50µL with serial dilutions of Fab’, DTFab’, DTFab’FITC/N_3_, or the unlabelled parental mAb (concentrations ranging from 150 µg.mL^-1^ to 1.83*10^−2^ µg.mL^-1^, i.e 3 µM to 3.66*10^−4^ µM for Fab’ fragments, and 1 µM to 1.22*10^−4^ µM for the mAb). After 10 min of incubation at 4°C, 50 µL of 1 µg.mL^-1^ labelled parental antibody was added to the wells. After 20 min of incubation at 4°C, the plate was washed twice, and AF647 fluorescence intensity was acquired by flow cytometry on a MACSQuant plate reader (Miltenyi Biotech).

### Antibody purification from hybridoma supernatant

Engineered hybridomas selected for antibody production were cultured in CELLine™ 1000 flasks (7340389, VWR) flasks following the manufacturer’s recommendations. Cells were harvested every other week, supernatant collected by centrifugation (90 g, 5 min), and density gradient purification (Lymphoprep™, 07861, Stem Cell Technologies) was performed to remove dead cells before reseeding them. Supernatant was filtered through a 0.2 μm filter, and supplemented with 10 mM imidazole (I2399, Sigma-Aldrich). Supernatant was incubated with 2 mL of pre-washed Ni-NTA beads (30210, Qiagen) at 4°C on a tube roller for 1h. Ni-NTA beads were transferred to a disposable chromatography column (7321010, Bio-Rad), washed with 5 column volumes (CV) of ice-cold wash buffer [50 mM Tris pH=8.0, 150 mM NaCl, 20 mM imidazole] and eluted using 2*3 mL of ice-cold elution buffer [50 mM Tris pH=8.0, 150 mM NaCl, 250 mM imidazole]. Buffer exchange to ice-cold PBS or sortase buffer [50 mM Tris pH=7.5, 150 mM NaCl] was performed via ultracentrifugation at 4°C with Amicon Ultra-15 centrifugal filter units (UFC901024, Merck-Millipore). Antibodies from WT hybridomas were purified from medium using Protein G GraviTrap columns (28-9852-55, Sigma-Aldrich). Columns were washed with 10 mL of binding buffer [0.02 M sodium phosphate, pH=7.0], 20 mL of sample per column was added, and washed with 15 mL of binding buffer. Collection tubes were pre-filled with 1mL of neutralizing buffer [1 M Tris-HCl, pH=9.0], and column was eluted using 3 mL of elution buffer [0.1 M Glycine-HCl, pH=2.7]. Antibodies were immediately dialysed to PBS or sortase buffer by ultracentrifugation at 4°C. Antibody absorbance was measured using a Nanodrop 2000 (ThermoFisher Scientific) and the UV-Vis program. If necessary, antibody absorbance was corrected by measuring the absorbance of the free label at 280 nm and λmax, and calculating the correction factor F= A_280label_/A_λmaxlabel_. Antibody absorbance was corrected by A_Ab_=A_280Ab_ – (A_λmaxAb_*F), and concentration was calculated by Beer-Lambert’s law, using a correction factor of 1.4 (parental mAb) or 1.35 (Fab’ fragment). Protein purity was assessed on a 12% SDS-PAGE using SYPRO Ruby Protein Gel Stain (S12000, Thermo Fisher Scientific).

### Sortase production and purification

eSrt(2A-9) and eSrt(4S-9) pET29b were a gift from David Liu (Addgene # 75145 and #75146 respectively). Sortase mutants were produced in BL21(DE3) *E. coli* as reported^27^. Briefly, chemically competent BL21(DE3) were transformed by heat shock, and grown overnight at 30°C in selective media. The next day, selective media was inoculated at OD_600_∼0.05, and bacteria were grown at 37°C, 220 rpm to an OD_600_∼0.6, and induced with 1 mM IPTG (I5502, Sigma-Aldrich) for 16h at 25°C. Bacteria were collected by centrifugation, the pellet washed with [50 mM Tris, 150 mM NaCl], and frozen at -20°C overnight. Pellets were thawed on ice, and resuspended in lysis buffer [50 mM Tris, 150 mM NaCl, 10 mM imidazole (I0250, Sigma-Aldrich), 20 µg.mL^-1^ of protease inhibitor cocktail (4693159001, Roche), 10% (v/v) glycerol]. Suspension was lysed by sonication on ice (3*30sec, 25% amplitude), and centrifuged (8600 g, 30 min, 4°C). Supernatant was collected, and protein was isolated using Ni-NTA beads. After 1h of incubation at 4°C, beads were washed with 100 CV of ice-cold wash buffer [50 mM Tris pH=7.5, 150 mM NaCl, 10 mM imidazole]. Protein was eluted using 2*4 CV ice-cold elution buffer [50 mM Tris pH=7.5, 150 mM NaCl, 500 mM imidazole, 10% (v/v) glycerol], and washed by ultracentrifugation at 4°C using a 3kDa filter (UFC900324, Merck-Millipore) to remove imidazole. Protein concentration was measured on Nanodrop 2000 (MW=17752 Da, ε_280nm_=14565 M^-1^cm^-1^), and sortase was stored at -80°C in sortase buffer [50 mM Tris pH=7.5, 150 mM NaCl] supplemented with 10%(v/v) glycerol. Protein purity was assessed on a 12% SDS-PAGE using SYPRO Ruby Protein Gel Stain (S12000, Thermo Fisher Scientific).

### Sortase-mediated transpeptidation

To assess sortase-mediated conjugation, small nucleophilic peptides [H-GGG-C-K(FAM)-NH_2_], [H-GGG-C-K(FITC)-NH_2_] and [H-GGG-K(N3)-NH_2_] were constructed via solid-phase peptide synthesis using a Fmoc/tBu approach and Rink amide resin to yield C-terminal amidated peptides. FAM and FITC were coupled to Lys side chain before peptide cleavage and purified on preparative HPLC using linear gradient of acetonitrile. Peptides were cleaved from the resin by treating the peptidyl-resin with a cleavage cocktail of TFA/TIS/H_2_O (95:2.5:2.5). [H-GGG-C-K(FAM)-NH_2_], [H-GGGCK(FITC)-NH_2_] and [H-GGGK(N3)-NH_2_] purity was determined by analytical HPLC. Antibodies stored in PBS were dialysed to sortase buffer via ultracentrifugation at 4°C. For small scale optimization reactions, 2 µg of Fab’ fragment was reacted, while 500 µg-1 mg was used for large scale reactions using proportional volumes. eSrt2A-9- and eSrt4S-9-mediated ligations were carried out using 0.75 equivalent of sortase and 50 equivalents of nucleophile per equivalent of Fab’ fragment (final antibody concentration 20 μM), in sortase buffer supplemented with CaCl_2_ at 100 mM final concentration. Reactions were carried out at 37°C for 1h. Small scale reactions were stopped by addition of 100 mM EDTA, and the volume corresponding to 250 ng of Fab’ fragment was loaded onto a reducing SDS-PAGE gel for analysis. Large scale reactions were incubated at 4°C on a rotating incubator with 100 μL of pre-washed Ni-NTA beads. After 30 min, the mixture was transferred to a disposable spin column, and the flow through collected. Beads were washed with 2*500 µL of wash buffer [50 mM Tris pH=8, 150 mM NaCl, 10 mM imidazole], and the resulting flow-throughs were pooled. Resulting volume was purified by fast protein liquid chromatography (NGC Quest, BioRad) on an ENrich™ SEC70 10×300 column (7801070, BioRad) in sortase buffer. Proteins were concentrated on a 0.5 mL 10 kDa filter (UFC501024, Merck-Millipore) by ultracentrifugation. Product purity was assessed on a reducing SDS-PAGE and analysed for fluorescence and protein signals using the SYPRO Ruby Protein Gel Stain (S12000, ThermoFisher) on a Typhoon Trio+ imager (GE Healthcare). In addition, the *m/z* of the conjugates was analysed on Bruker Microflex LRF MALDI-TOF equipment. Samples were analyzed within a concentration range of 0.3-0.5 mg.mL^-1^ in MilliQ using 1 µL each of matrix-sample-matrix sown on the MALDI plate. Matrix: Sinapic acid (trans-3,5-dimethoxy-4-hydroxycinnamic acid) from Sigma Aldrich (MERCK, D7927) at 10 mg.mL^-1^ in H2O/MeCN (1/1) + 0.1 % TFA.

## IV. Results

### Generation of a genetically engineered cell line secreting Fab’ fragments

To produce Fab’ fragments suitable for dual site-specific labelling we sought to alter the immunoglobulin domain within the genome of an anti-hCD20-producing hybridoma cell line, which expresses a mAb of the mIgG1 isotype with a κ LC (Cκ). We hypothesized this could be achieved by adapting our recently developed CRISPR/HDR approach^23^, in which we genetically engineered the rat IgG2a IgH locus, for modification of the murine IgH and IgK loci. We started with the genetic modification of the HC (figure 1A and B). To this end, we selected a guide RNA (gRNA-H.m1) that directs the Cas9 protein to the CH1 region of the mIgG1 IgH locus^28^. The HDR template consists of ∼600bp 5’ and 3’ homology arms (HA) flanking the intended modification site in the hinge region of the HC. It was designed to insert a modified hinge sequence, a GGGGS linker, the LAETGG tag for chemoenzymatic modification with eSrt2A-9, the hexahistidine-tag (His-tag) and a stop codon. Additionally, the HDR template contained elements to confer blasticidin resistance upon genomic integration (supplementary table S1). The parental cell line anti-CD20 WT was transfected with the gRNA-H.m1 and HDR template, and subjected to blasticidin selection pressure. Following antibiotic selection, we isolated DNA from the bulk population to confirm the insert integration in resistant cells by PCR. Using a primer that hybridizes directly upstream of the 5’HA, and one specific for the insert (IRES) (figure 1B), we could detect a unique band at the expected height of 788bp (figure 1C), attesting for successful insertion of the HDR template into the targeted locus.

**Figure 1.**
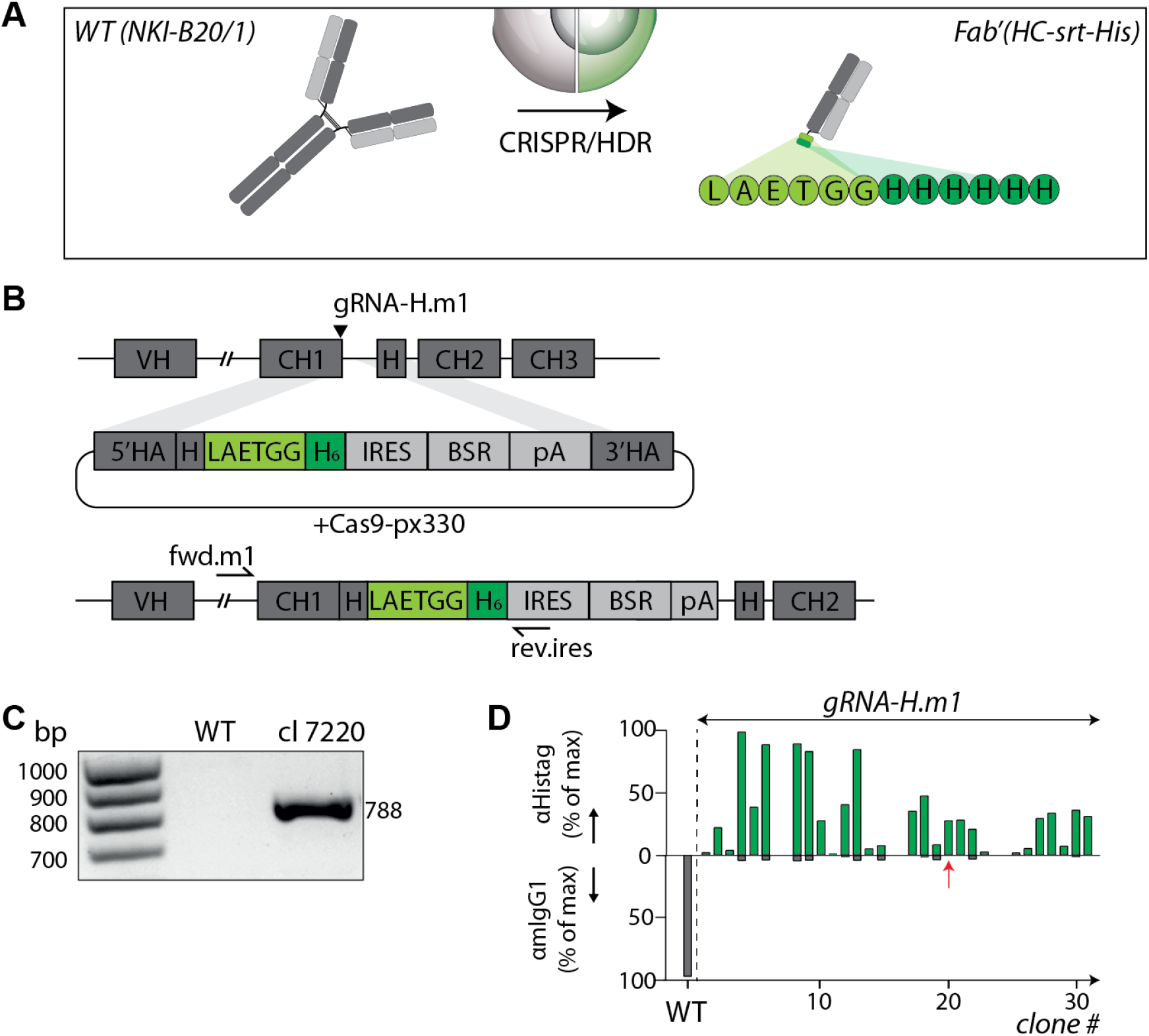
Generation of Fab’ fragment-secreting cell lines from parental hybridoma NKI-B20/1. **A**. and **B**. General strategy for CRISPR/Cas9 editing. gRNA-H.m1 cuts in the CH1 domain to introduce the first five residues of the hinge region followed by a GGGGS semi-flexible linker, a LAETGG sortag motif and a polyHis-tag (HHHHHH). **C**. Genomic PCR on DNA isolated from the bulk population after blasticidin selection shows that resistant cells have integrated the insert. **D**. Mean fluorescence intensity (MFI) of anti-His-tag and anti-mIgG1 on BJAB cells following incubation with antibody-containing supernatants collected from single cell clones shows that most clones have been successfully edited. The red arrow indicates the clone selected for Fab’ production and subsequent genome editing.

We performed limiting dilution to generate monoclonal cell lines, and after 14 days of culture we harvested the supernatant of single cell clones, and used it to stain hCD20-expressing BJAB cells (a human Burkitt lymphoma B cell line). To assess genetic engineering, we determined whether the clones were secreting wild type mAbs or His-tagged Fab’ fragments using secondary antibodies against mIgG1 and His-tag. Supernatant of over 60% of single cell colonies that grew out after antibiotic selection were positive for the His-tag, and negative for mIgG1 signal (figure 1D), indicating that these edited monoclonal cell lines did not secrete the WT mAb anymore, but instead produced a His-tagged Fab’ fragment. We selected a high producing monoclonal cell line based on His-tag signal intensity, cell growth rate, and strictly negative mIgG1 signal to eliminate potential contamination by the WT cell line (clone 7220, Fab’(HC-srt-his)), and expanded it to pursue the second step of genome editing.

### Generation of a genetically engineered cell line secreting DTFab’ fragments

We continued with the second step of genome editing by genetically modifying the LC to obtain a DTFab’-producing cell line. To edit the LC (figure 2A), we designed an HDR template consisting of ∼600bp 5’ and 3’ HA flanking the C-terminus of the κ LC (Cκ), and in-frame inserted an LPESGG sortag for chemoenzymatic modification with eSrt4S-9, and a C-terminal c-Myc epitope tag (Myc-tag, EQKLISEEDL) for screening purposes (supplementary table S2). We additionally inserted a puromycin resistance gene (figure 2B). We selected a guide RNA (gRNA-L.mκ) directing Cas9 to the C-terminus of the LC, and transfected the anti-CD20 Fab’(HC-srt-his)-secreting hybridoma. After puromycin selection, we collected resistant cells and genomic PCR confirmed insert integration in the Cκ region in this bulk population (figure 2C). Monoclonal cell lines were obtained by limiting dilution, after which we proceeded with flow cytometry screening using the monoclonal supernatants to stain hCD20-expressing BJAB cells and to analyse His-tag and Myc-tag signals. Only dual tagged Fab’ will be equipped with both the His-tag and Myc-tag. Interestingly, most cell lines secreted high levels of Fab’ fragments harbouring both His-tag and Myc-tag, while others had overall low production levels (figure 2D). Importantly, none of the monoclonal cell lines were producing antibody fragments displaying only a His-tag, indicating the robustness of our strategy. A monoclonal cell line (clone7220×4887, DTFab’ (LC-srt-Myc)) was selected based on DTFab’ expression level, and expanded for further characterization.

**Figure 2.**
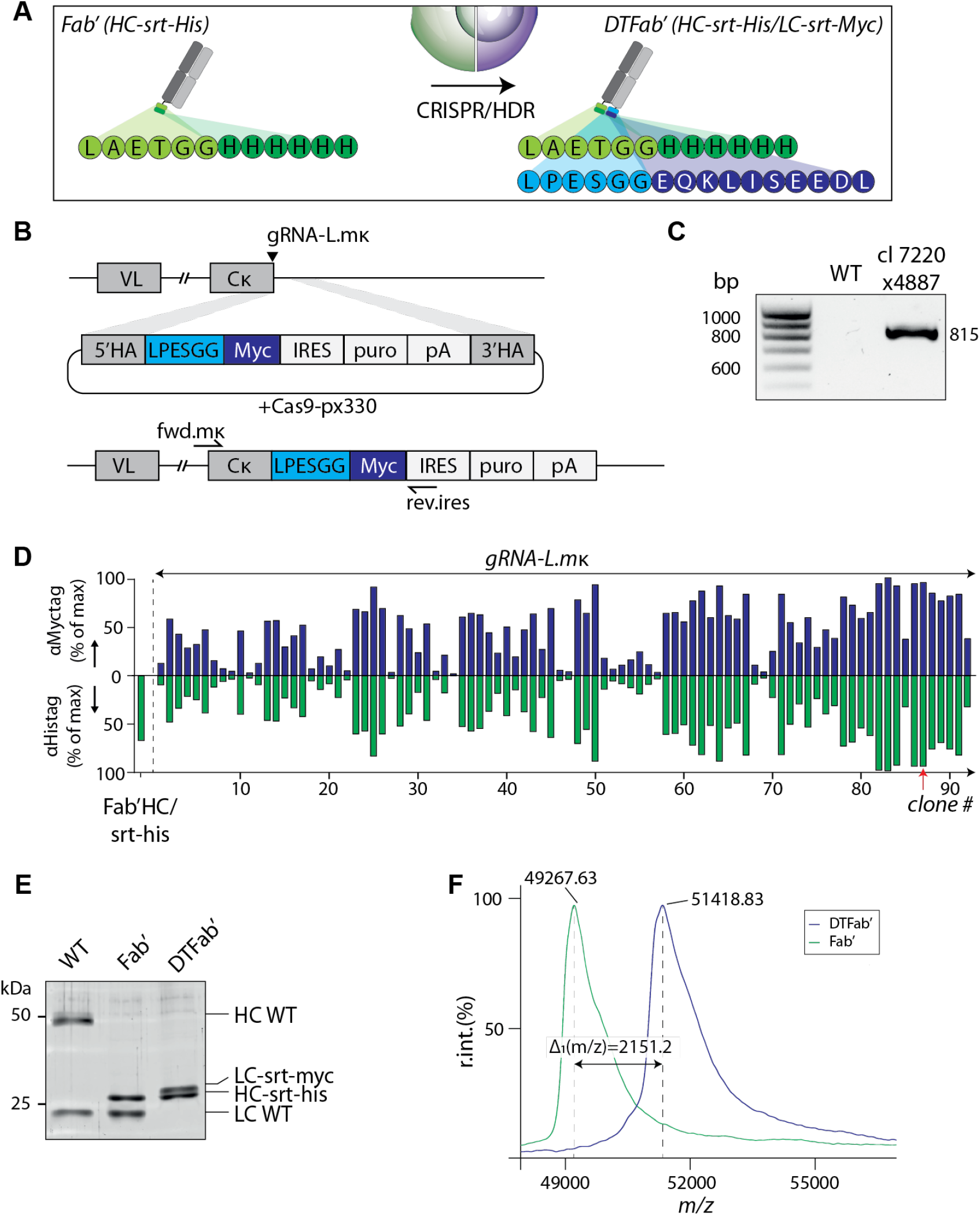
Generation of DTFab’ fragment-secreting cell lines from Fab’ modified hybridoma. **A**. and **B**. General strategy for CRISPR/Cas9 editing. gRNA-L.mκ cuts in the κ light chain to introduce a GGGGS semi-flexible linker, a LPESGG sortag motif and a Myc-tag (EQKLISEEDL). **C**. Genomic PCR performed on the bulk population after puromycin selection shows that resistant cells have integrated the insert. **D**. Mean fluorescence intensity (MFI) of His-tag and anti-Myctag on BJAB cells following incubation with antibody-containing supernatants collected from single cell shows near perfect overlap of signals coming from the heavy and light chains for each clone, attesting for a high efficacy of the selection strategy. The red arrow indicates the clone selected for DTFab’ production. **E**. SDS-PAGE visualisation of WT, Fab’ and DTFab’ proteins. **F**. MALDI-TOF analysis of Fab’ and DTFab’ shows engineering of the LC leads to expected increase of molecular weight, Δ_1_=2151.2 Da, expected 2059.2 Da.

### Engineered antibodies are functionalized site-specifically and orthogonally

Having the parental hybridoma and the two engineered daughter cell lines in hand (WT, Fab’(HC-srt-his) and DTFab’ (HC-srt-his, LC-srt-Myc), respectively), we cultured the cells for antibody production, and purified products from the supernatants. Subsequently we analysed the products by sodium dodecyl sulfate polyacrylamide gel electrophoresis (SDS-PAGE), which confirmed that genomic engineering of the HC resulted in an expected shift from 50kD for the WT HC to approximately 25kDa for the Fab’(HC-srt-his). Genetic editing of the Igκ locus resulted in an increase of the molecular weight of the LC, in line with the insertion of the LPESGG motif and Myc-tag, as seen by SDS-PAGE and MALDI-TOF analysis (figure 2E,2F and S2A). We then proceeded to confirm the orthogonality of the two sortags. The functionality and specificity of the two different sortags were examined by incubating the DTFab’ with either eSrt2A-9 or eSrt4S-9, and an excess of 6-carboxyfluorescein (FAM)-containing substrate peptide **1** (GGGCK(FAM)) (figure 3A). The resulting products were analysed by fluorescent SDS-PAGE (figure 3B and S1A). A fluorescent (488nm (FAM)) protein band was detected, which overlapped with the HC when DTFab’ was reacted in presence of eSrt2A-9, and conversely, the LC was fluorescently labelled when in presence of eSrt4S-9. This confirms that the sortag motifs are functional, and that both HC and LC could be functionalized orthogonally with the respective sortase mutants without detectable cross-reactivity.

**Figure 3.**
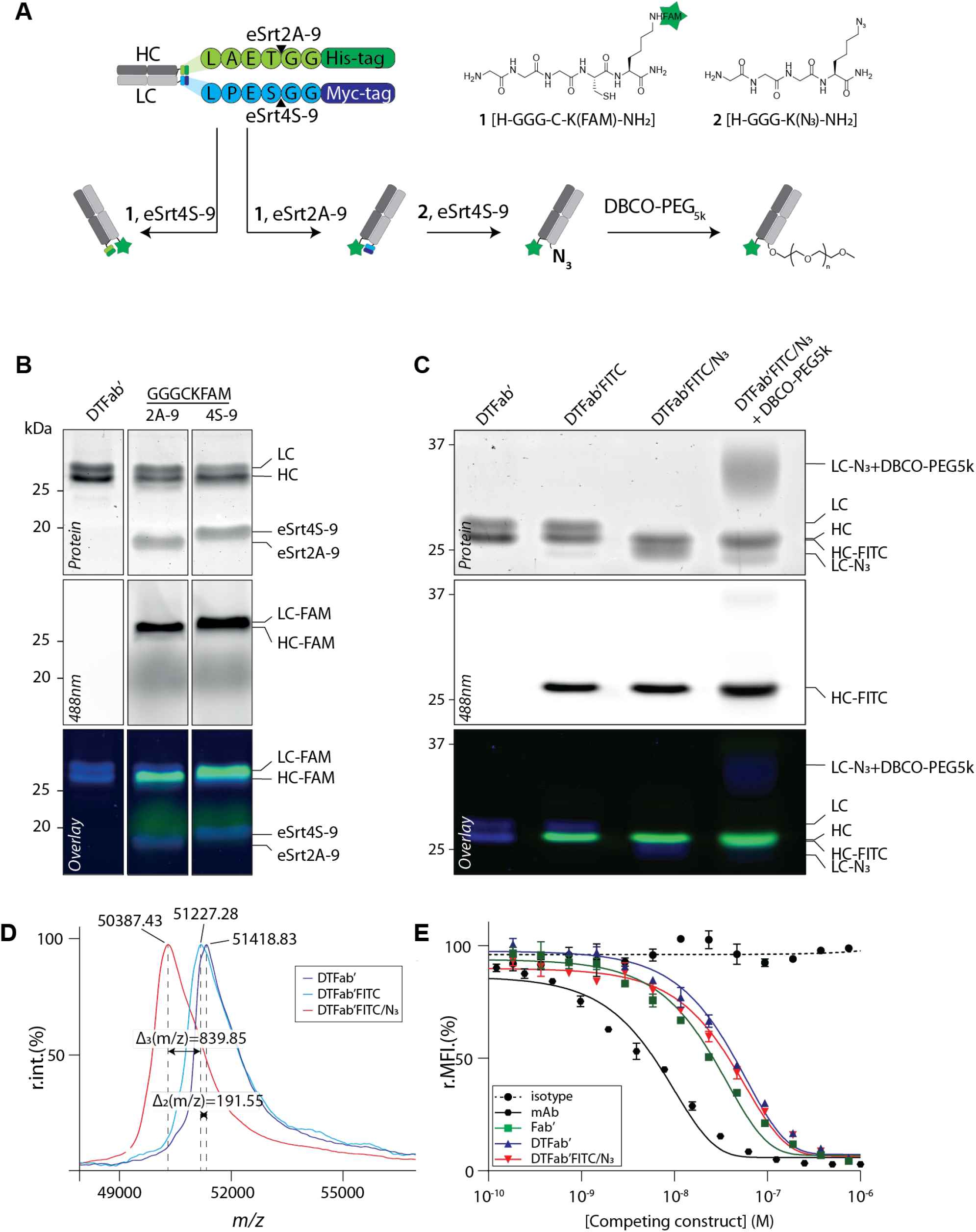
Efficient dual site-specific labelling of DTFab’ onto both HC and LC. **A**. General strategy and peptides used for labelling. **B**. SDS PAGE and fluorescence (488nm) analysis of sortagging reactions performed separately on the HC (eSrt2A-9) or LC (eSrt4S-9) confirms specificity of the enzymes for their respective cleavage sites. **C**. SDS PAGE analysis of DTFab’ modified with GGG-C-K(FITC)-NH_2_ on the HC, and GGG-K(N_3_)-NH_2_ on the LC confirms introduction of two distinct cargos site-specifically onto the engineered Fab’. **D**. MALDI-TOF analysis of DTFab’, DTFab’FITC and DTFab’FITC/N3 shows that proteins undergo the expected change in molecular weight during chemoenzymatic ligation (Δ_2_=191.55 Da, expected 249.21 Da; Δ_3_=839.85 Da, expected 975.06 Da). **E**. Antigen binding competition assay of each engineered protein against mAb-AF647 reveals that proteins do not lose binding affinity to their target following CRISPR/Cas9 editing and sequential dual site-specific labelling.

### Dual site-specific labelling of DTFab’ does not hamper its binding affinity

As a proof-of-concept of dual site-specific modification, we proceeded to sequential sortagging to introduce two different payloads on the DTFab’ (figure 3A). We first incubated DTFab’ with eSrt2A-9 and an excess of fluorescent peptide **1** (GGG-C-K(FITC)-NH_2_), followed by protein purification (near quantitative conversion, isolated yield slightly above 60%). The second functionality was introduced by incubation with eSrtA4S-9 and an excess of the azide-functionalized peptide **2** (GGG-K(N_3_)-NH_2_) and subsequent protein purification (near quantitative conversion, isolated yield slightly above 50%). All antibody fragments were analysed by (fluorescent) SDS-PAGE (figure 3C) and MALDI-TOF (figure 3D and S2B). As observed by SDS-PAGE analysis, the FITC signal overlaps only with the HC, while ligation of the azido peptide onto the LC leads to a complete shift to a product with a reduced molecular weight, attesting for site-specific incorporation of the cargo. In addition, strain promoted alkyne-azide cycloaddition (SPAAC) of the dual functionalized Fab’ with an excess of PEG_5k_-DBCO leads to a selective increase in size of the LC, showing the functionality of the azide for further diversification of the antibody fragment. MALDI-TOF analysis of the products before and after site-specific modification of the HC and LC confirmed mass changes corresponding to the respective transformations. Finally, to confirm that genome editing and dual site-specific payload conjugation did not affect antigen binding, we performed a competitive antigen binding assay against the AF647-labelled parental antibody on BJAB cells. Analysis of the AF647 signal indicates that all constructs compete with the parental antibody for CD20 binding (figures 3E and S3B). Importantly, while genetic alteration from a mAb to a Fab’ fragment leads to an expected decrease in avidity (divalent to monovalent binding), neither the introduction of a tag at the C-terminus of the LC, nor sequential introduction of two payloads at the C-termini of the HC and LC significantly altered their binding ability. Taken together, our data shows that we can engineer a hybridoma cell line to produce high titers of a Fab’ fragment bearing two distinct chemoenzymatic modification sites on the HC and LC, allowing for orthogonal dual site-specific conjugation of different functionalities.

## V. Discussion

In this study, we present for the first time a strategy to generate dual-tagged Fab’ fragments bearing two orthogonal site-specific modification tags (DTFab’), which can readily be utilized for the chemoenzymatic conjugation with two distinct cargos. Extending on our previous work^23^, we demonstrate the successful engineering of a hybridoma secreting mIgG1 antibodies (anti-CD20 WT) to a stable daughter cell line producing Fab’ fragments carrying an eSrt2A-9 (LAETGG) motif on its HC, and an eSrt4S-9 (LPESGG) motif on its κ LC (anti-CD20 DTFab’). This method enables robust biorthogonal engineering of virtually any antibody-secreting hybridoma to reproducibly produce high yields of modified DTFab’ molecules.

The anti-CD20 DTFab’ daughter cell line secreted high titers of the engineered protein (>1.5mg/mL), which could easily be isolated and modified by orthogonal sortase-mediated transpeptidation. Upon incubation of the DTFab’ protein with either sortase mutant eSrt2A-9 or eSrt4S-9 in the presence of a fluorescent substrate peptide, we detected exclusive fluorescent labelling of the HC or LC, respectively. As expected^24^, the presence of two sortase recognition sites (LAETGG and LPESGG) in close proximity did not affect the specificity of eSrt2A-9 and eSrt4S-9 for their respective motifs, thus enabling site-specific conjugation of two distinct payloads at the C-termini of the HC and LC. As a proof-of-concept, we introduced a fluorescent dye on the HC, and a reactive organic azide on the LC by sequential sortagging (DTFab’FITC/N_3_). MALDI-TOF and (fluorescent) SDS-PAGE analysis confirmed the identity and the purity of the final product. Importantly, we demonstrated that the target binding capacity of such obtained dual labelled Fab’ fragment is uncompromised, as anticipated due to very mild reaction and purification conditions, and the maximized distance between the cargos and the binding site of the Fab’ fragment.

ADC generation through site-specific conjugation offers the advantage to obtain highly homogenous products with a uniform DAR, thus increasing the therapeutic window^11^. Several strategies to introduce drugs site-specifically have been developed^29^, notably via engineered cysteine introduction at the carboxyl-terminus or the Fab’domain^30^, unnatural amino acids, glycans, or short peptide tags catalysed by enzymes^22^. Among these, sortagging offers the advantage of near-quantitative conversion yields, fast reaction times, and high versatility in the payload, since most cargo can easily be functionalized with an amino-terminal polyglycine motif and thus act as sortase substrate. Others have demonstrated efficient generation of ADCs using transiently expressed recombinant antibodies bearing sortase A LPXTG motifs^31^. Our straightforward CRISPR/HDR hybridoma engineering platform offers the advantage of generating continuously expressing stable cell lines, resulting in an unlimited source of site-specific functionalizable antibody fragments. More recently, several groups have reported disulphide bond reduction and re-bridging strategies to modify native proteins^32–35^. While elegant, such strategies could lead to affinity loss following re-bridging and influence antibody rigidity and flexibility^36,37^, which has recently been linked to therapeutic efficacy^38^. By targeting the C-termini of both HC and LC, we do not expect that the conjugation strategy applied in our approach will affect structure or flexibility of the antibody fragment, which is supported by the fact the target binding capacity was uncompromised.

Others have described strategies to introduce two cargos site-specifically on a single protein by introducing a combination of tags recognized by orthogonal enzymes, self-labelling tags, engineered residues or inteins^39–44^. All these strategies have their advantages, but share the requirement to know the sequence of the antibody’s variable region to accommodate recombinant expression. Our strategy relies on targeting of the antibody’s constant domains and is therefore in principle applicable to any hybridoma, without the need to sequence the variable region. This can be achieved by simply adapting the gRNAs in the Cas9 plasmid, and the homology arms in the HDR template to the constant domain sequence of the isotype and species of choice.

Current challenges faced by anti-tumour ADCs in the clinic include tumour infiltration, target downregulation by tumour cells, and acquired drug resistance^11^. It is estimated that less than 2% of administrated dose of ADC reaches the tumour^20,45^, the large size of the conjugates being one of the major hurdles. Fab’ fragments represent an attractive alternative to mAbs for generating ADCs because of their smaller size, which could lead to a better tumour infiltration. Our platform can be used to target multiple distinct cytotoxic agents to tumour cells. The Goldie-Coldman model^46^ stipulates that an early treatment is the best way to preventing evolution of acquired drug resistance. Accordingly, combination of several chemotherapeutic drugs to limit the chances of tumour resistance is a powerful strategy^47,48^, but doses and combinations are limiting due to an invariably associated higher toxicity. Over the past 30 years of ADC development^49–52^, it has been well-documented that ADCs offer the advantage of improved therapeutic window, and reduced off-target toxicity. We foresee that harnessing our platform to generate Fab’ molecules, equipped with two highly cytotoxic payloads with distinct mechanisms of action to target tumours, could lead to a better therapeutic outcome, while reducing the toxicity of the combination therapy. In efforts to explore various combinations of drugs, synthetically easily accessible, rapid and versatile platform such as the one presented in this work is instrumental. The versatility can additionally be harnessed to introduce other entities in addition to a drug, such as PEG chains to tune the half-life of the conjugates, and fluorophores or radioligands for theragnostic purposes.

## Author contributions

J.M.S.v.d.S., M.V. and F.A.S. conceived the project. K.M.B., C.G.F., F.A.S. and M.V. provided guidance and support. C.M.L., J.M.S.v.d.S., and M.V. designed experiments. C.M.L., J.M.S.v.d.S. and I.R.T., performed the experiments. M.P.K., F.J.v.D., Z.W., L.S., D.v.D. and A.C., contributed experimentally. C.M.L., F.A.S. and M.V. wrote the paper with assistance from all authors. All authors reviewed the manuscript.

## Data and materials availability

All data needed to evaluate the conclusions are present in the paper and/or the Supplementary Materials. The described plasmids used in this study are deposited in plasmid repository of Addgene (www.addgene.org/), or may be requested from the authors.

## List of supplementary material

Figure S1 – SDS-PAGE analysis of DTFab’

Figure S2 – MALDI-TOF analysis of the conjugates, full spectra

Figure S3-Flow cytometry analysis

Supplementary table 1 –Ighg1 locus and Fab’ HDR template sequence.

Supplementary table 2 –Cκ locus and DTFab’ HDR template sequence.

