## Supplementary data for "Dual site-specific chemoenzymatic antibody fragment conjugation using CRISPR-based hybridoma engineering"

**Supplementary information**

**Figure S1 – SDS-PAGE analysis of DTFab’**

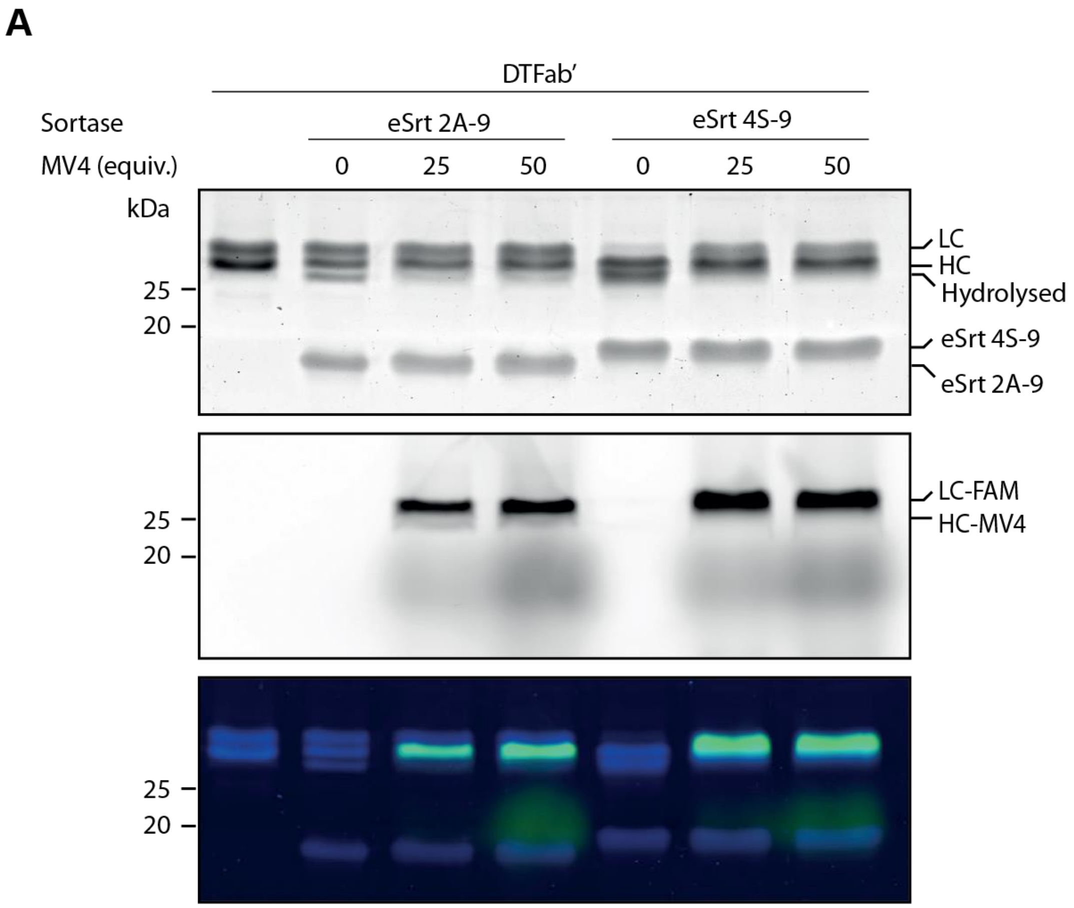


**Figure S1 – SDS-PAGE analysis of DTFab’. A.** SDS PAGE and fluorescence (488nm) analysis of sortagging reactions performed separately on the HC (eSrt2A-9) or LC (eSrt4S-9) with different equivalents of MV4[GGG-C-K(FAM)-$NH_2$] (0, 25, 50) confirms specificity of the enzymes for their respective cleavage sites.

**Figure S2 – MALDI-TOF analysis of the conjugates, full spectra**

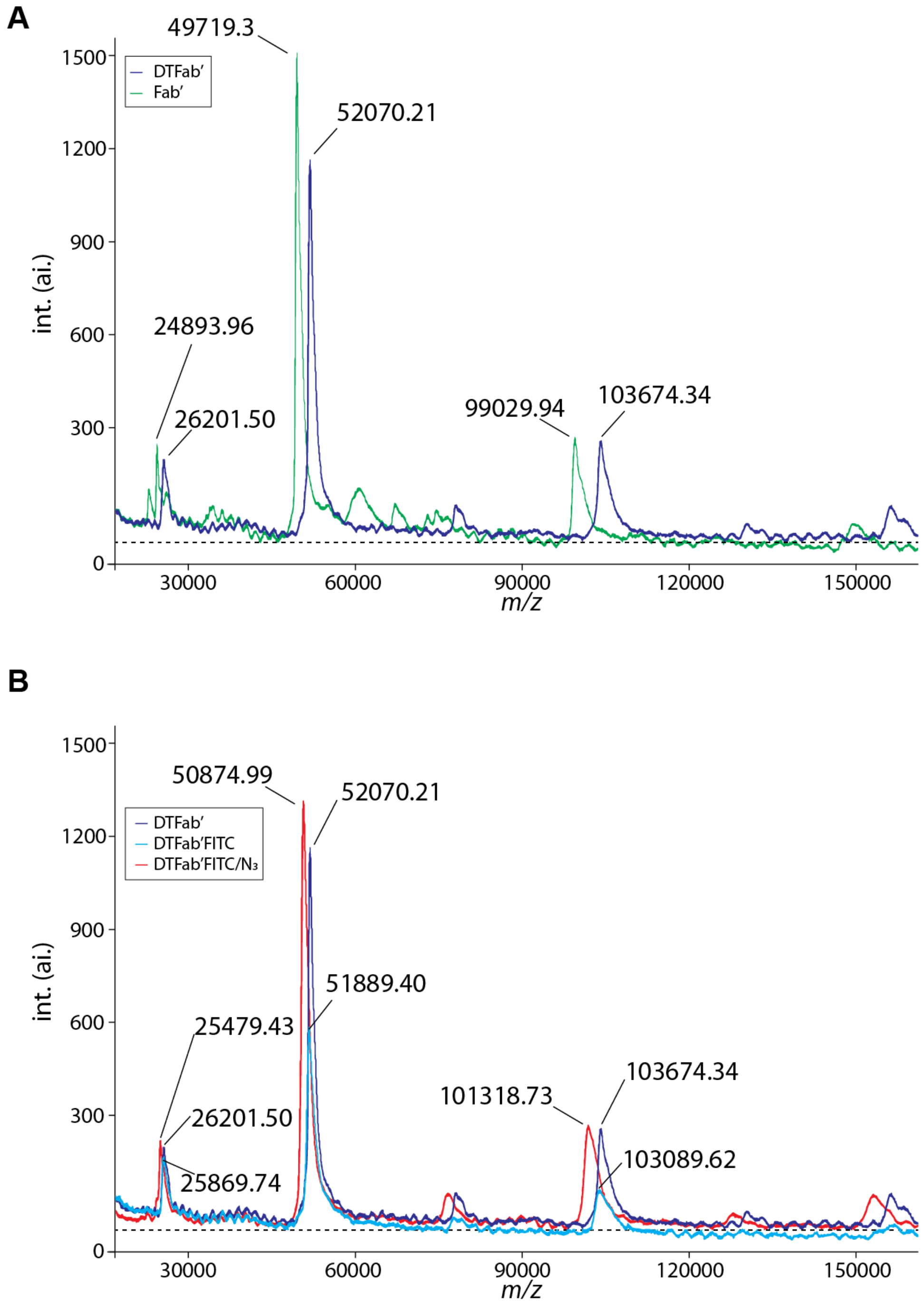


**Figure S2 – MALDI-TOF analysis of the conjugates, full spectra**. **A.** Full spectra (20-160kDa) of Fab' vs DTFab' and **B.** Full spectra (20-160kDa) of DTFab' vs DTFab'FITC vs DTFab'FITC/$N_3$ acquired by MALDI-TOF, non-normalized intensities.

**Figure S3- Flow cytometry analysis**

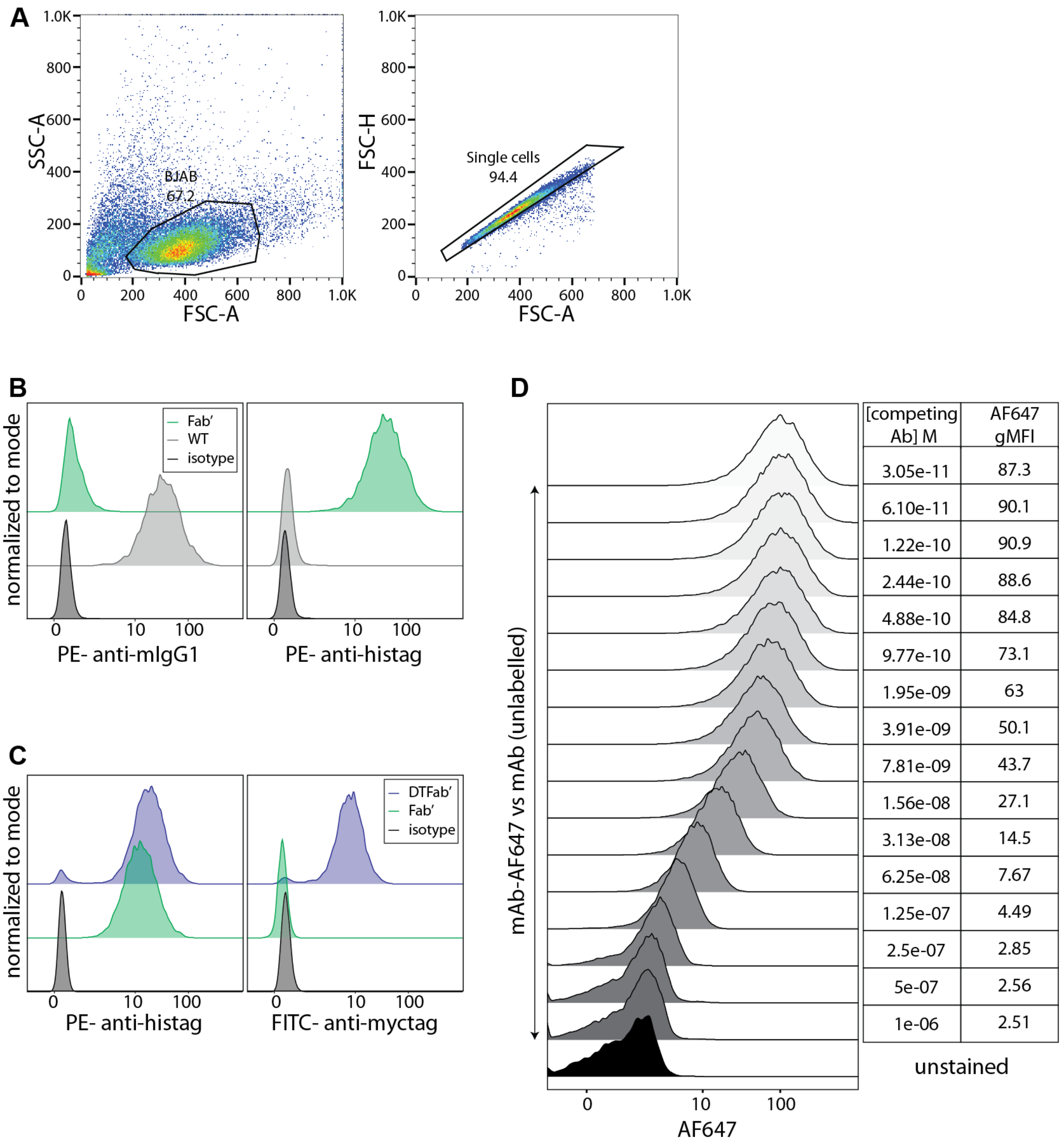


**Figure S3- Flow cytometry analysis**. **A.** Gating of BJAB used in flow cytometry binding experiments. **B.** Example of flow cytometry raw data obtained during Fab' screening. mIgG1 and His-tag signals measured on BJAB cells after incubation with culture supernatants derived from WT hybridoma (grey) and the selected Fab'clone (green) generated during the first genome editing step. **C.** Example of flow cytometry raw data obtained during DTFab' screening. His-tag and Myc-tag signals measured on BJAB cells after incubation with culture supernatants derived from Fab' hybridoma (green) and the selected DTFab'clone (blue) generated during the second genome editing step. **D.** Example of flow cytometry raw data obtained during antigen binding competition assay (here for mAb vs mAb-AF647). AF647 MFI quantification for serial dilutions of competing unlabelled antibody.

**Supplementary table 1 –Ighg1 locus and Fab' HDR template sequence. A.** Ighg1 sequence used to design the HDR template. Coding regions (CH1 and hinge) are highlighted in bold. PAM sequence is highlighted in red, and gRNA-H.m1 annealing site is underlined. **B.** HDR template sequence.

| **Table 1A** | **Sequence** |
|---|---|
| Ighg1_DNA (+ strand) ENSMUSG00000076614 | CAGACAGCTGGGGGGCGGGGGTGAACACAGATACCCATACTGGAAAGCAG<br>GTGGGGCATTTTCCTAGGAACGGGACTGGGCTCAATGGCCTCAGGTCTCAT<br>CTGGTCTGGTGATCCTGACATTGATAGGCCCAAATGTTGGATATCACCTAC<br>TCCATGTAGAGAGTCGGGGACATGGGAAGGGTGCAAAAGAGCGGCCTTCT<br>AGAAGGTTTGGTCCTGTCCTGTCCTGTCTGACAGTGTAATCACATATACTTT<br>TTCTTGTAG**CCAAAACGACACCCCCATCTGTCTATCCACTGGCCCCTGG<br>ATCTGCTGCCCAAACTAACTCCATGGTGACCCTGGGATGCCTGGTCAA<br>GGGCTATTTCCCTGAGCCAGTGACAGTGACCTGGAACTCTGGATCCCT<br>GTCCAGCGGTGTGCACACCTTCCCAGCTGTCCTGCAGTCTGACCTCTA<br>CACTCTGAGCAGCTCAGTGACTGTCCCCTCCAGCACCTGGCCCAGCCA<br>GACCGTCACCTGCAACGTTG*CCC*ACCCGGCCAGCAGCACCAAGGTGGA<br>CAAGAAAATTG**GTGAGAGGACGTATAGGGAGGAGGGGTTCACTAGAGGT<br>GAGGCTCAAGCCATTAGCCTGCCTAAACCAACCAGGCTGGACAGCCATCA<br>CCAGGAAATGGATCTCAGCCCAGAAGATCGAAAGTTGTTCTTCTCCCTTCT<br>GGAGATTTCTATGTCCTTTACACTCATTGGTTAATATCCTGGGTTGGATTCC<br>CACACATCTTGACAAACAGAGACAATTGAGTATCACCAGCCAAAAGTCAT<br>ACCCAAAAACAGCCTGGCATGACCTCACACCAGACTCAAACTTACCCTACC<br>TTTATCCTGGTGGCTTCTCATCTCCAGACCCCAGTAACACATAGCTTTCTCT<br>CCACAG**TGCCCAGGGATTGTGGTTGTAAGCCTTGCATATGTACAG**GTA<br>AGTCAGTAGGCCTTTCACCCTGACCCCAGATGCAACAAGTGGCCATGTTAG<br>AGGGTGGCCCAGGTATTGACCTATTTCCACCTTTCTTCTTCATCCTTAGTCC<br>CAGAAGTATCATCTGTCTTCATCTTCCCCCCAAAGCCCAAGGATGTGCTCA<br>CCATTACTCTGACTCCTAAGGTCACGTGTGTTGTGGTAGACATCAGCAAGG<br>ATGATCCCGAGGTCCAGTTCAGCTGGTTTGTAGATGATGTGGAGGTGCACA<br>CAGCTCAG |
| Ighg1_AA P01868 | **AKTTPPSVYPLAPGSAAQTNSMVTLGCLVKGYFPEPVTVTWNSGSLSSGV<br>HTFPAVLQSDLYTLSSSVTVPSSPRPSETVTCNVAHPASSTKVDKKIVPRDC<br>GCKPCICT** |

| **Table 1B** | **Sequence** |
|---|---|
| 5'HA | CAGTATTGTCCAGATTGTGTGCAGCCATATGGCCCAGGTATAAGAGGTTTA<br>ACAGTGGAACACAGATGCCCACATCAGACAGCTGGGGGGCGGGGGTGAAC<br>ACAGATACCCATACTGGAAAGCAGGTGGGGCATTTTCCTAGGAACGGGAC<br>TGGGCTCAATGGCCTCAGGTCTCATCTGGTCTGGTGATCCTGACATTGATA<br>GGCCCAAATGTTGGATATCACCTACTCCATGTAGAGAGTCGGGGACATGG<br>GAAGGGTGCAAAAGAGCGGCCTTCTAGAAGGTTTGGTCCTGTCCTGTCCTG<br>TCTGACAGTGTAATCACATATACTTTTTCTTGTAGCCAAAACGACACCCCC<br>ATCTGTCTATCCACTGGCCCCTGGATCTGCTGCCCAAACTAACTCCATGGT<br>GACCCTGGGATGCCTGGTCAAGGGCTATTTCCCTGAGCCAGTGACAGTGAC<br>CTGGAACTCTGGATCCCTGTCCAGCGGTGTGCACACCTTCCCAGCTGTCCT<br>GCAGTCTGACCTCTACACTCTGAGCAGCTCAGTGACTGTCCCCTCCAGCAC |

| | |
|---|---|
| | CTGGCCCAGCCAGACCGTCACCTGCAACGTTGCTCATCCGGCATGTAGCACCAAGGTAGACAAAAAGATA |
| Modified hinge | GTGCCCAGGGATTGT |
| MCS | GGATCCCTCGAG |
| G4S-LAETGG-His-tag-stop | GGAGGTGGCGGGTCACTTGCTGAGACAGGGGGACACCATCACCATCACCATTGA |
| MCS | CAATTGCATATGACCGGTGAGCTC |
| IRES | TCCCTCCCCCCCCCCTAACGTTACTGGCCGAAGCCGCTTGGAATAAGGCCGGTGTGCGTTTGTCTATATGTTATTTTCCACCATATTGCCGTCTTTTGGCAATGTGAGGGCCCGGAAACCTGGCCCTGTCTTCTTGACGAGCATTCCTAGGGGTCTTTCCCCTCTCGCCAAAGGAATGCAAGGTCTGTTGAATGTCGTGAAGGAAGCAGTTCCTCTGGAAGCTTCTTGAAGACAAACAACGTCTGTAGCGACCCTTTGCAGGCAGCGGAACCCCCCACCTGGCGACAGGTGCCTCTGCGGCCAAAAGCCACGTGTATAAGATACACCTGCAAAGGCGGCACAACCCCAGTGCCACGTTGTGAGTTGGATAGTTGTGGAAAGAGTCAAATGGCTCTCCTCAAGCGTATTCAACAAGGGGCTGAAGGATGCCCAGAAGGTACCCCATTGTATGGGATCTGATCTGGGGCCTCGGTGCACATGCTTTACATGTGTTTAGTCGAGGTTAAAAAACGTCTAGGCCCCCCGAACCACGGGGACGTGGTTTTCCTTTGAAAAACACGATGATAA |
| MCS | TCTAGAGTCGACGTTAAC |
| Blasticidin resistance | ATGAAGCCTTTGTCTCAAGAAGAATCCACCCTCATTGAAAGAGCAACGGCTACAATCAACAGCATCCCCATCTCTGAAGACTACAGCGTCGCCAGCGCAGCTCTCTCTAGCGACGGCCGCATCTTCACTGGTGTCAATGTATATCATTTTACTGGGGGACCTTGTGCAGAACTCGTGGTGCTGGGCACTGCTGCTGCTGCGGCAGCTGGCAACCTGACTTGTATCGTCGCGATCGGAAATGAGAACAGGGGCATCTTGAGCCCCTGCGGACGGTGCCGACAGGTGCTTCTCGATCTGCATCCTGGGATCAAAGCCATAGTGAAGGACAGTGATGGACAGCCGACGGCAGTTGGGATTCGTGAATTGCTGCCCTCTGGTTATGTGTGGGAGGGCTAA |
| MCS | GAGCTCGCTAGC |
| SV40 polyA | CTGTGCCTTCTAGTTGCCAGCCATCTGTTGTTTGCCCCTCCCCCGTGCCTTCCTTGACCCTGGAAGGTGCCACTCCCACTGTCCTTTCCTAATAAAATGAGGAAATTGCATCGCATTGTCTGAGTAGGTGTCATTCTATTCTGGGGGGTGGGGTGGGGCAGGACAGCAAGGGGGAGGATTGGGAAGACAATAGCAGGCATGCTGGGGATGCGGTGGGCTCTATGG |
| MCS | AGATCTTTAATTAA |
| 3’HA | GGTGAGAGGACGTATAGGGAGGAGGGGTTCACTAGAGGTGAGGCTCAAGCCATTAGCCTGCCTAAACCAACCAGGCTGGACAGCCATCACCAGGAAATGGATCTCAGCCCAGAAGATCGAAAGTTGTTCTTCTCCCTTCTGGAGATTTCTATGTCCTTTACACTCATTGGTTAATATCCTGGGTTGGATTCCCACACATCTTGACAAACAGAGACAATTGAGTATCACCAGCCAAAAGTCATACCCAAAAACAGCCTGGCATGACCTCACACCAGACTCAAACTTACCCTACCTTTATCCTGGTGGCTTCTCATCTCCAGACCCCAGTAACACATAGCTTTCTCTCCACAGTGCCCAGGGATTGTGGTTGTAAGCCTTGCATATGTACAGGTAAGTCAGTAGGCCTTTCACCCTGACCCCAGATGCAACAAGTGGCCATGTTAGAGGGTGGCCCAGGTATTGACCTATTTCCACCTTTCTTCTTCATCCTTAGTCCCAGAAGTATCATCTGTCTTCATCTTCCCCCCAAAGCCCAAGGATGTGCTCACCATTACTCTGACTCCTAAGGTCACGTGTGTTGTGGTAGACATCAGCAAGGATGATCCCGAGGTCCAGTTCAGCTGGTTTGT |

**Supplementary table 2 –Cκ locus and DTFab' HDR template sequence. A.** Cκ sequence used to design the HDR template. Coding regions (CH1 and hinge) are highlighted in bold. PAM sequence is highlighted in red, and gRNA-L.Cκ annealing site is underlined. **B.** HDR template sequence.

| **Table 2A** | **Sequence** |
|---|---|
| Ck_DNA (+ strand)<br><br>ENSMUSG00000076609 | TCTGTCTGAAGCATGGAACTGAAAAGAATGTAGTTTCAGGGAAGAAAGGC<br>AATAGAAGGAAGCCTGAGAATATCTTCAAAGGGTCAGACTCAATTTACTTT<br>CTAAAGAAGTAGCTAGGAACTAGGGAATAACTTAGAAACAACAAGATTGT<br>ATATATGTGCATCCTGGCCCCATTGTTCCTTATCTGTAGGGATAAGCGTGCT<br>TTTTTGTGTGTCTGTATATAACATAACTGTTTACACATAATACACTGAAATG<br>GAGCCCTTCCTTGTTACTTCATACCATCCTCTGTGCTTCCTTCCTCAG**GGGC**<br>**TGATGCTGCACCAACTGTATCCATCTTCCCACCATCCAGTGAGCAGTT**<br>**AACATCTGGAGGTGCCTCAGTCGTGTGCTTCTTGAACAACTTCTACCC**<br>**CAAAGACATCAATGTCAAGTGGAAGATTGATGGCAGTGAACGACAAAA**<br>**TGGCGTCCTGAACAGTTGGACTGATCAGGACAGCAAAGACAGCACCTA**<br>**CAGCATGAGCAGCACCCTCACGTTGACCAAGGACGAGTATGAACGAC**<br>**ATAACAGCTATACCTGTGAGGCCACTCACAAGACATCAACTTCACCCA**<br>**TTGTCAAGAGCTTCAACAGGAATGAGT**GTTAGAGACAA***AGG***TCCTGAGA<br>CGCCACCACCAGCTCCCCAGCTCCATCCTATCTTCCCTTCTAAGGTCTTGGA<br>GGCTTCCCCACAAGCGACCTACCACTGTTGCGGTGCTCCAAACCTCCTCCC<br>CACCTCCTTCTCCTCCTCCTCCCTTTCCTTGGCTTTTATCATGCTAATATTTG<br>CAGAAAATATTCAATAAAGTGAGTCTTTGCACTTGAGATCTCTGTCTTTCTT<br>ACTAAATGGTAGTAATCAGTTGTTTTTCCAGTTACCTGGGTTTCTCTTCTAA<br>AGAAGTTAAATGTTTAGTTGCCCTGAAATCCACCACACTTAAAGGATAAAT<br>AAAACCCTCCACTTGCCCTGGTTGGCTGTCCACTACATGGCAGTCCTTTCTA<br>AGGTTCACGAGTACTATTCATGGCTTATTTCTCTGGGCCATGGTAGGTTTGA<br>GGAGGCATACTTCCTAGTTTTCTTCCCCTAAGTCGTCAAAGTCCTGAAGGG<br>GGACAGTCTTTACAAGCACATGTTCTGTAATCTGATTCAACCTACCCAGTA<br>AACTTGGCGAAGCAAAGTAGAATCATTATCACAGGAAGCAAAGGCAACCT<br>AAATGTGCA |
| Ck_AA P01837 | **RADAAPTVSIFPPSSEQLTSGGASVVCFLNNFYPKDINVKWKIDGSERQNG**<br>**VLNSWTDQDSKDSTYSMSSTLTLTKDEYERHNSYTCEATHKTSTSPIVKSF**<br>**NRNEC** |

| **Table 2B** | **Sequence** |
|---|---|
| 5'HA | TCTGTCTGAAGCATGGAACTGAAAAGAATGTAGTTTCAGGGAAGAAAGGC<br>AATAGAAGGAAGCCTGAGAATATCTTCAAAGGGTCAGACTCAATTTACTTT<br>CTAAAGAAGTAGCTAGGAACTAGGGAATAACTTAGAAACAACAAGATTGT<br>ATATATGTGCATCCTGGCCCCATTGTTCCTTATCTGTAGGGATAAGCGTGCT<br>TTTTTGTGTGTCTGTATATAACATAACTGTTTACACATAATACACTGAAATG<br>GAGCCCTTCCTTGTTACTTCATACCATCCTCTGTGCTTCCTTCCTCAGGGGC<br>TGATGCTGCACCAACTGTATCCATCTTCCCACCATCCAGTGAGCAGTTAAC<br>ATCTGGAGGTGCCTCAGTCGTGTGCTTCTTGAACAACTTCTACCCCAAAGA<br>CATCAATGTCAAGTGGAAGATTGATGGCAGTGAACGACAAAATGGCGTCC<br>TGAACAGTTGGACTGATCAGGACAGCAAAGACAGCACCTACAGCATGAGC<br>AGCACCCTCACGTTGACCAAGGACGAGTATGAACGACATAACAGCTATAC |

| | |
|---|---|
| | CTGTGAGGCCACTCACAAGACATCAACTTCACCGATTGTAAAGAGTTTTAA<br>TAGCAATGAATGCGGATCC |
| MCS | CTCGAG |
| G4S-LPESGG-<br>Myc-tag-STOP | GGAGGTGGCGGGTCATTGCCTGAATCCGGTGGAGAGCAGAAGCTGATCTC<br>AGAGGAGGACCTGTGA |
| MCS | CAATTGCATATGACCGGTGAGCTC |
| IRES | TCCCTCCCCCCCCCCTAACGTTACTGGCCGAAGCCGCTTGGAATAAGGCCG<br>GTGTGCGTTTGTCTATATGTTATTTTCCACCATATTGCCGTCTTTTGGCAAT<br>GTGAGGGCCCGGAAACCTGGCCCTGTCTTCTTGACGAGCATTCCTAGGGGT<br>CTTTCCCCTCTCGCCAAAGGAATGCAAGGTCTGTTGAATGTCGTGAAGGAA<br>GCAGTTCCTCTGGAAGCTTCTTGAAGACAAACAACGTCTGTAGCGACCCTT<br>TGCAGGCAGCGGAACCCCCCACCTGGCGACAGGTGCCTCTGCGGCCAAAA<br>GCCACGTGTATAAGATACACCTGCAAAGGCGGCACAACCCCAGTGCCACG<br>TTGTGAGTTGGATAGTTGTGGAAAGAGTCAAATGGCTCTCCTCAAGCGTAT<br>TCAACAAGGGGCTGAAGGATGCCCAGAAGGTACCCCATTGTATGGGATCT<br>GATCTGGGGCCTCGGTGCACATGCTTTACATGTGTTTAGTCGAGGTTAAAA<br>AACGTCTAGGCCCCCCGAACCACGGGGACGTGGTTTTCCTTTGAAAAACAC<br>GATGATAA |
| MCS | TCTAGAGTCGACGTTAAC |
| Puromycin<br>resistance | ATGACCGAGTACAAGCCCACAGTGCGGCTGGCCACAAGAGATGATGTGCC<br>TAGAGCTGTGCGGACACTGGCCGCTGCTTTCGCTGATTACCCTGCCACCAG<br>ACACACTGTGGACCCCGACAGACACATCGAGAGAGTGACCGAGCTGCAAG<br>AGCTGTTCCTGACCAGAGTCGGCCTGGACATCGGCAAAGTGTGGGTTGCAG<br>ATGATGGCGCCGCTGTGGCTGTGTGGACAACACCTGAATCTGTGGAAGCCG<br>GCGCTGTGTTCGCTGAGATCGGACCTAGAATGGCCGAGCTGTCTGGCAGCA<br>GACTGGCTGCTCAGCAGCAGATGGAAGGACTGCTGGCTCCCCACAGACCT<br>AAAGAGCCTGCCTGGTTCCTGGCTACCGTGGGAGTGTCTCCTGACCACCAA<br>GGCAAAGGCCTGGGATCTGCTGTTGTGCTGCCTGGCGTTGAGGCTGCTGAA<br>AGAGCTGGCGTCCCAGCCTTCCTGGAAACAAGCGCCCCTAGAAACCTGCCT<br>TTCTACGAGAGACTGGGCTTCACCGTGACCGCCGATGTGGAAGTTCCTGAG<br>GGACCAAGAACCTGGTGCATGACCAGAAAGCCTGGCGCCTAA |
| MCS | GAGCTCGCTAGC |
| SV40 polyA | CTGTGCCTTCTAGTTGCCAGCCATCTGTTGTTTGCCCCTCCCCCGTGCCTTC<br>CTTGACCCTGGAAGGTGCCACTCCCACTGTCCTTTCCTAATAAAATGAGGA<br>AATTGCATCGCATTGTCTGAGTAGGTGTCATTCTATTCTGGGGGGTGGGGT<br>GGGGCAGGACAGCAAGGGGGAGGATTGGGAAGACAATAGCAGGCATGCT<br>GGGGATGCGGTGGGCTCTATGG |
| MCS | AGATCTTTAATTAA |
| 3’HA | TGAAGACAAAGGTCCTGAGACGCCACCACCAGCTCCCCAGCTCCATCCTAT<br>CTTCCCTTCTAAGGTCTTGGAGGCTTCCCCACAAGCGACCTACCACTGTTGC<br>GGTGCTCCAAACCTCCTCCCCACCTCCTTCTCCTCCTCCTCCCTTTCCTTGGC<br>TTTTATCATGCTAATATTTGCAGAAAATATTCAATAAAGTGAGTCTTTGCAC<br>TTGAGATCTCTGTCTTTCTTACTAAATGGTAGTAATCAGTTGTTTTTCCAGT<br>TACCTGGGTTTCTCTTCTAAAGAAGTTAAATGTTTAGTTGCCCTGAAATCCA<br>CCACACTTAAAGGATAAATAAAACCCTCCACTTGCCCTGGTTGGCTGTCCA<br>CTACATGGCAGTCCTTTCTAAGGTTCACGAGTACTATTCATGGCTTATTTCT<br>CTGGGCCATGGTAGGTTTGAGGAGGCATACTTCCTAGTTTTCTTCCCCTAA<br>GTCGTCAAAGTCCTGAAGGGGGACAGTCTTTACAAGCACATGTTCTGTAAT<br>CTGATTCAACCTACCCAGTAAACTTGGCGAAGCAAAGTAGAATCATTATCA<br>CAGGAAGCAAAGGCAACCTAAATGTGCA |
